# Investigations on Antarctic fish IgM drives the generation of an engineered mAb by CRISPR/Cas9

**DOI:** 10.1101/2023.10.04.560855

**Authors:** Alessia Ametrano, Bruno Miranda, Rosalba Moretta, Principia Dardano, Luca De Stefano, Umberto Oreste, Maria Rosaria Coscia

**Affiliations:** Institute of Biochemistry and Cell Biology, National Research Council of Italy, Naples, Italy; Institute of Applied Sciences and Intelligent Systems, National Research Council of Italy, Naples, Italy; Sanidrink s.r.l., Naples, Italy

## Abstract

IgM is the major circulating Ig isotype in teleost fish, showing in Antarctic fish unique features such as an extraordinary long hinge region, which plays a crucial role in antibody structure and function. In this work, we describe the replacement of the hinge region of a murine monoclonal antibody (mAb) with the peculiar hinge from Antarctic fish IgM. We use the CRISPR/Cas9 system as a powerful tool for generating the engineered mAb. Then, we assessed its functionality by using an innovative plasmonic substrate based on bimetallic nanoislands (AgAuNIs). The affinity constant of the modified mAb was 2.5-fold higher than the one obtained from wild-type mAb against the specific antigen. Here, we show the suitability of the CRISPR/Cas9 method for modifying a precise region in immunoglobulin gene loci. The overall results could open a frontier in further structural modifications of mAbs for biomedical and diagnostic purposes.

## Introduction

Antibody engineering represents a powerful technology for high-affinity peptides/proteins discovery, receptor binding, and epitope identification, or for investigating protein-protein interactions. In this context, hybridoma technology is considered a significant milestone in the history of monoclonal antibodies (mAbs) generation, which has revolutionized biomedical research (Köhler & Milstein, 1975). Nowadays, mAbs can be produced not only by hybridoma cells but also through a variety of expression systems, such as yeasts, bacteria, mammalian cells, or phages (Frenzel *et al*, 2013). Although ensuring the production of large amounts of antibodies, all these methods have some disadvantages, such as random integration of transgenes, unbalanced production of heavy (H) and light (L) chains, time-consuming procedures, and high costs. These issues have been partly overcome through the use of the genetic engineering approach. Early approaches to targeted genomic modifications of hybridomas were applied to renew antibody production in deficient immunoglobulin (Ig) mutant cell line (Baker *et al*, 1988) or to replace IgH constant region gene locus from mouse to human (Fell *et al*, 1989). However, these traditional genetic manipulations tend to be ineffective and require multistep selectable markers, *e*.*g*., neomycin resistance cassette.

At the end of 90s an innovative genetic engineering approach, known as genome editing, has revolutionized the idea to modify the genomes of organisms (Woolf, 1998). It allows a gene sequence to be inserted, deleted, or modified in a precise manner at a targeted gene locus, only inducing DNA double-strand breaks (DSBs) that stimulate error-prone non-homologous end joining (NHEJ) or homology-directed repair (HDR). The first approaches of genome editing were carried out through the nucleases with reprogrammable targeting specificity, such as zinc finger nucleases (ZFNs) (Urnov *et al*, 2005) or transcription-activator-like effector nucleases (TALENs), which require the design and generation of a nuclease pair for every genomic target (Doudna & Sontheimer, 2014). The Clustered Regularly Interspaced Short Palindromic Repeats (CRISPR)-Cas9 (CRISPR-associated protein 9) system has led to a revolution in genome editing applications (Doudna & Charpentier, 2014). Interestingly, the CRISPR-Cas9 system was discovered as a prokaryotic adaptive immune system (Ishino *et al*, 1987; Hermans *et al*, 1991; Mojica & Montoliu, 2016) and over the past decade it has been widely employed to perform genome editing in mammalian cells (Ran *et al*, 2013).

Given several potential advantages, such as the capacity to use multiple guide RNAs to target multiple genes simultaneously in the same cell with high efficiency and specificity (Cong *et al*, 2013), this technology has been successfully applied also in the field of immunology. Some examples come from the use of the CRISPR/Cas9 technology to edit mouse and human IgH gene loci for obtaining class switch recombination or the secretion of fragment antigen-binding (Fab) fragments, without enzymatic digestion to remove the fragment crystallizable (Fc) region (Cheong *et al*, 2016). Another proof of the great versatility of this technology is the modification of the variable region of mAbs. This allows for changing the antigenic specificity of hybridoma cells rapidly (Pogson *et al*, 2016). CRISPR/Cas9 has been also applied to correct major histocompatibility complex (MHC) mismatches in cell transplantation (Kelton *et al*, 2017). Very recently, the CRISPR/Cas9 system has been used to produce site-specifically modified antibody conjugates as therapeutic and diagnostic tools (Khoshnejad *et al*, 2018) or to obtain isotype-switched chimeric antibodies (van der Shoot *et al*, 2019). Overall, several studies demonstrated that the manipulation of the Fc region increases antibody effector functions that may be exploited for different applications, such as antiviral or anti-cancer treatment (Wang, 2018; Liu, 2020; Jin *et al*, 2022). However, the employment of the CRISPR/Cas9 system for engineering the hinge region has yet to be reported and only a few works describing hinge modifications by site-direct mutagenesis are currently available (Dell’Acqua *et al*, 2006; Yan *et al*, 2012; Ashoor *et al*, 2018; Suzuki *et al*, 2018). The hinge region is crucial in the antibody architecture, linking Fc to the Fab arms and conferring the segmental flexibility, required for the recognition and binding of a wide range of antigenic epitopes (Adlersberg, 1976; Tan *et al*, 1990). However, it lacks homology to any Ig constant region domain. Hinge is distinguished by unique features, not seen in any other region of the Ig molecule: i) location in the central part of the heavy chains, between the two Fab arms and the Fc region. ii) a highly variable amino acid composition, particularly rich in cysteines and prolines: cysteines form inter-chain inter-changeable disulfide bonds; prolines confer conformational flexibility that allows waving, rotation of Fab arms, and wagging of the Fc fragment, thus facilitating antigen binding and triggering of effector functions (Carayannipoulos & Capra, 1993; Ho *et al*, 2005). It has been demonstrated that changes in length or of some key amino acid residues that account for the hinge flexibility and structural characteristics, can improve the modulation of the effector functions of human IgG1 (Dell’Acqua *et al*, 2006) or the fragmentation resistance of Fc-fused bispecific antibodies (Suzuki *et al*, 2018).

One of the most uncommon hinges of vertebrate Ig is that exclusively found in Antarctic fishes: IgM H chain has a remarkably long hinge region, ranging from 14 to 22 amino acid residues, highly polymorphic, and rich in prolines and glycines (Coscia *et al*, 2000; Coscia *et al*, 2011; Coscia *et al*, 2012). Given these peculiar features, the hinge region has been suggested to provide the cold Ig molecule with higher flexibility to exert its function at very low kinetic energy in the Antarctic environment.

In this work, the structural peculiarities uncovered for cold-adapted Igs combined with the innovative application of the CRISPR-Cas9 system inspired the idea to generate an “antarctized” mAb (thereafter named anta-mAb). It was achieved by inserting the IgM hinge region sequence from an Antarctic fish species into the mouse *IgG1 H* chain constant region gene. Then, the impact of this structural modification on the performance of the engineered mAb was assessed by comparing it with its wild-type (WT) counterpart. This comparison was performed by immobilizing both mAbs on a bimetallic plasmonic nanoislands array and measuring their relative affinity for the target antigen.

## Results

### Knock-out of the *IgH* gene locus in the 9E10 hybridoma cell line

The experimental design consisted in targeting the *IgH* gene locus of the 9E10 murine hybridoma cell line, secreting mAbs specific for human c-Myc tag, by the CRISPR/Cas9 system (Fig 1A; Fig EV1).

**Figure 1.**
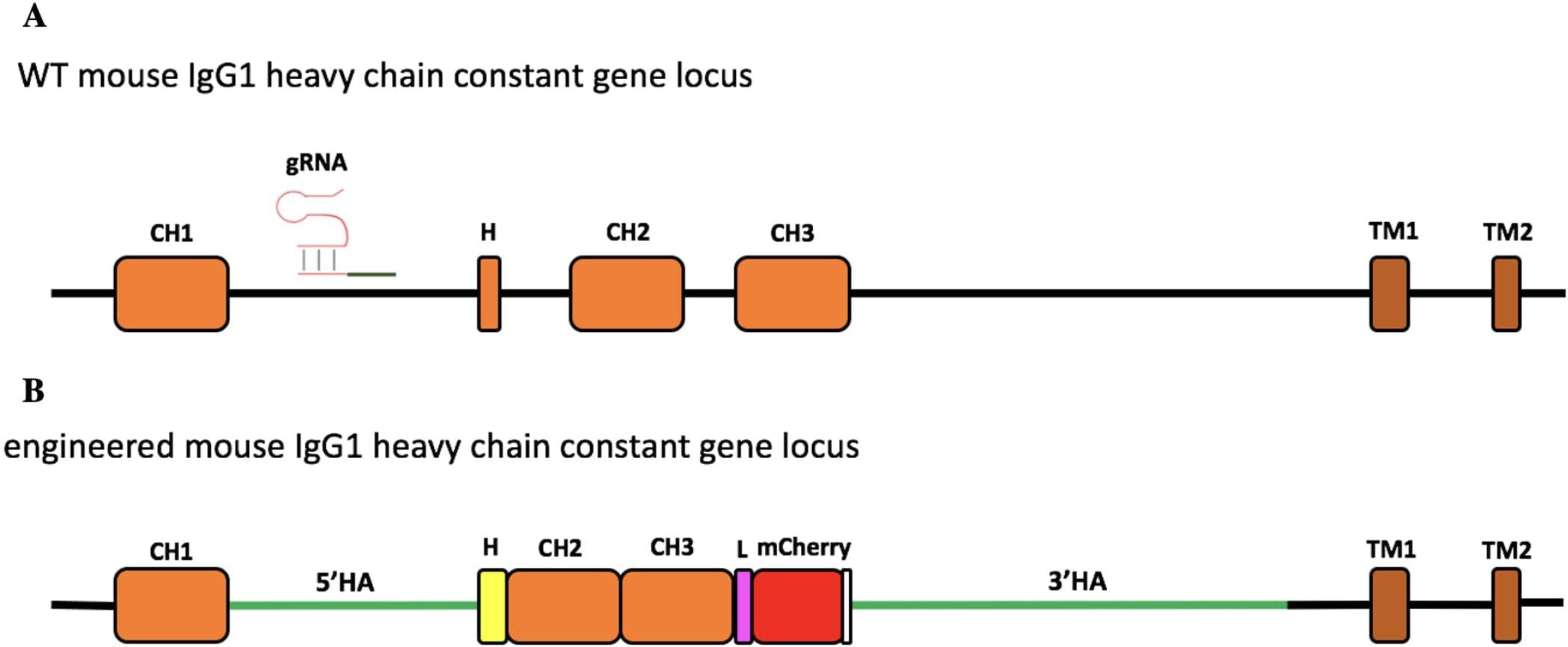
Schematic representation of engineering of the murine IgG1 heavy chain gene locus. A The region of the wild-type (WT) mouse IgG1 heavy chain constant gene locus involved in genome editing. The gRNA targeting the intron between the CH1 and hinge exon (H) is indicated. B The engineered IgH locus resulting from HDR-based exchange of the H sequence of mouse IgG1 with that of Antarctic fish IgM. The coding sequence of the insert contains the Antarctic fish hinge exon in yellow, followed by mouse IgG1 CH2 and CH3 exons, linked through a linker sequence (L) to mCherry, used as a selection marker. The stop codon is reported in white. The donor construct is flanked by the homology arms (5’HA and 3’HA, green lines) of 1005 and 2421 bp, respectively.

In the first step, the hinge exon and the region spanning the *CH2* and *CH3* exons were deleted. For this purpose, the gRNAs, essential for efficient cleavage and short indels of the mouse IgG1 H chain constant gene target, were designed to avoid the inactivation of the *IgH*-coding gene caused by frameshifts. Thus, to analyze the target specificity of gRNAs within the genomic context, E-CRISP was chosen as the optimal bioinformatics tool since it provides a flexible output and experiment-oriented design parameters, for designing and evaluating gRNAs (Heigwer, 2014; Doench *et al*, 2014; Xu *et al*, 2015). The input sequence used to design gRNAs was an annotated intronic region of mouse *IgG1 H* chain gene locus retrieved from Genbank NCBI (Fig EV1). Of seven gRNAs obtained, the two ones (gRNA1 and gRNA2) showing the best activity prediction score and the ideal location (153-bp and 41-bp upstream of the target site, respectively) were chosen for genome editing (Table EV1).

The two gRNAs were then cloned each into the CRISPR/Cas9 plasmid pX458, which enables the detection of Cas9 expression through the green fluorescence protein (GFP) (Ran *et al*, 2013) (Fig EV2). The sequencing confirmed that gRNAs were properly cloned into the pX458 plasmid (Fig EV3).

To perform gene knock-out, CRISPR plasmids, each containing gRNA1 or gRNA2, were transfected into the 9E10 hybridoma cells using the electroporation protocol. At 24 h post-transfection, GFP-positive cells were isolated by FACS for gRNA1 (6.8%, Fig 2D) and gRNA2 (22.0%, Fig 2H), relative to untreated cells (negative control), although the cell viability of the total cell population was ranging from 44.6% to 48.7% (Fig 2B and F).

**Figure 2.**
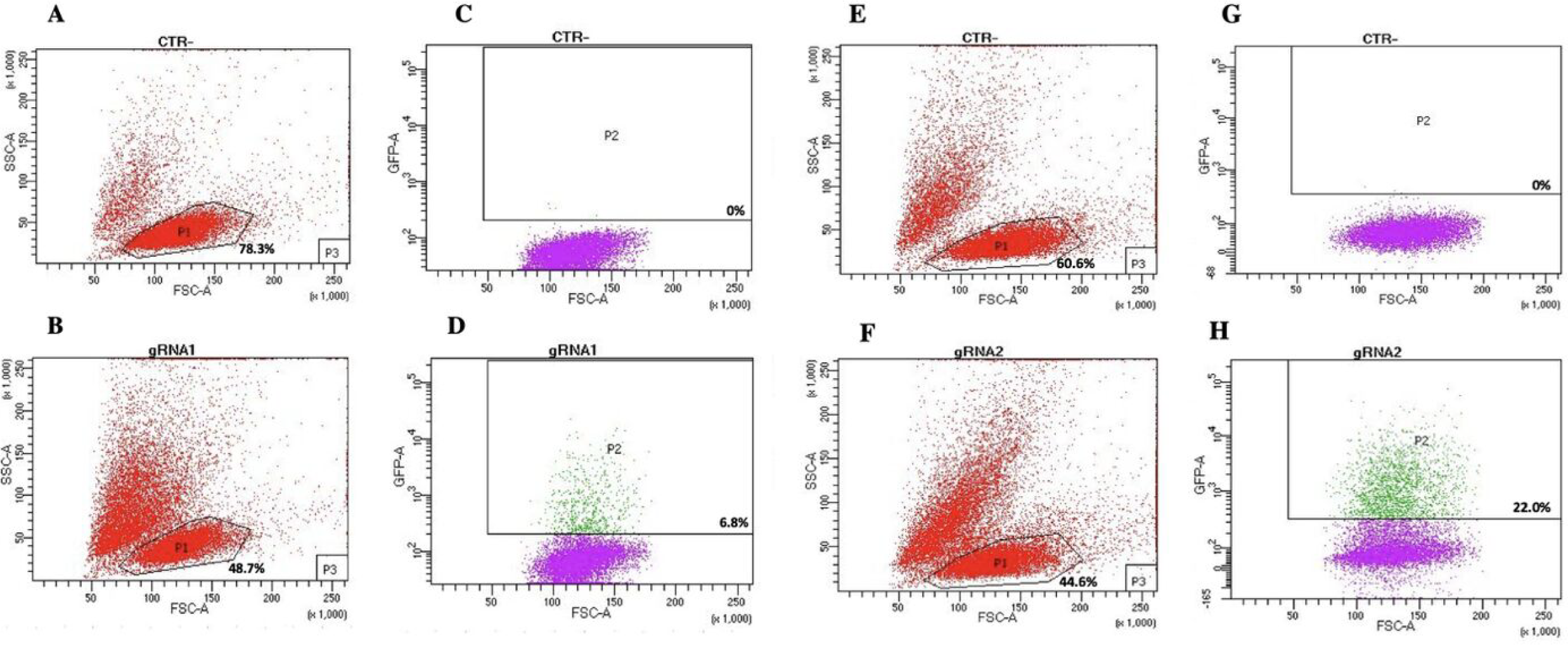
Flow cytometry analysis of gRNA1 and gRNA2 targeting of mouse *IgG1 H* chain gene locus in 9E10 hybridoma cells. A–H Flow cytometry dot plots show the cell viability (A, B, E, F) and Cas9 expression via 2A-GFP (C, D, G, H) in 9E10 hybridoma cells after electroporation with CRISPR plasmids pX458 containing gRNA1 (D) and gRNA2 (H) in comparison with untreated cells (C and G; CTR-). The percentage of GFP-positive cells among transfected cells is reported.

The effectiveness of Cas9 targeting in the mouse IgH genomic region was then assessed by PCR amplification, using primers that flanked the region of interest (Table EV2). The PCR products were cloned and then sequenced. Sequence analysis of two representative clones for gRNA1 and three for gRNA2 confirmed the expected Cas9-mediated deletion at the target site of the mouse IgG1 H chain constant gene locus (Fig 3A and B). In particular, mIgGgRNA2.2 and mIgGgRNA2.3 clones showed a 6- and 10-bp deleted region, respectively (Fig 3B), in line with the NHEJ mechanism preferentially adopted by cells for repairing DSBs (Mao *et al*, 2008).

**Figure 3.**
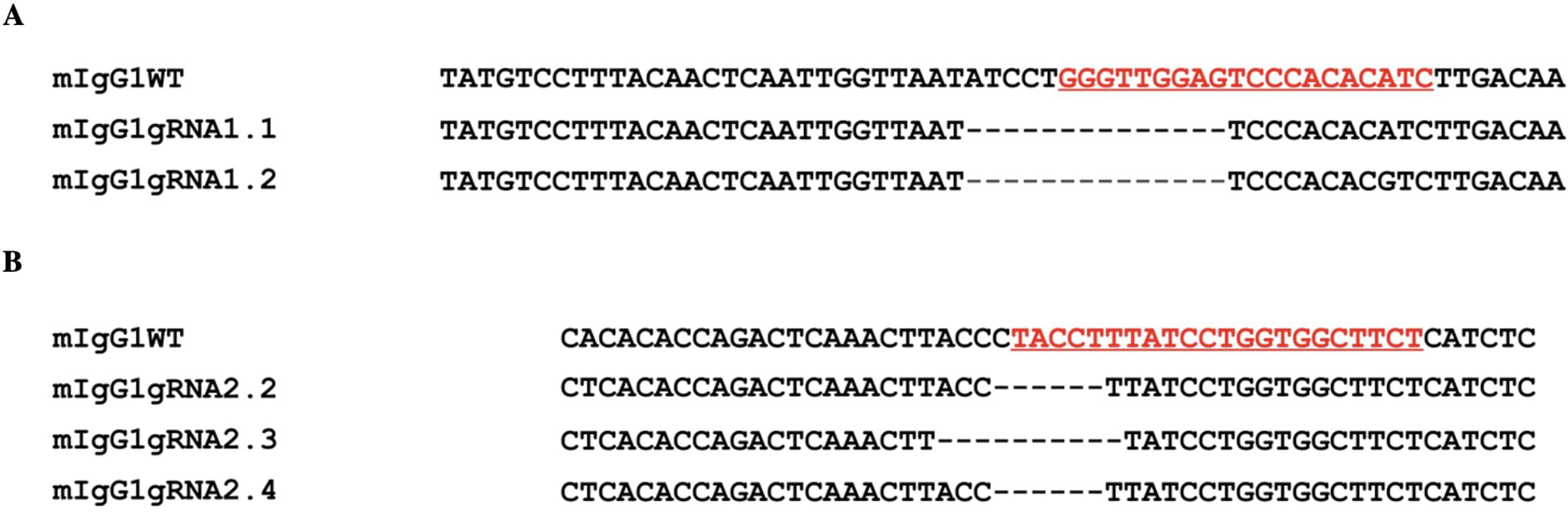
Evaluation of the Cas9 activity by PCR amplification of the targeted intron between the *CH1* and the hinge exon. A, B Nucleotide sequences from representative clones confirmed the expected Cas9-mediated deletion through gRNA1 (A; mIgG1gRNA1.1 and mIgG1gRNA1.2) and gRNA2 (B; mIgG1gRNA2.2, mIgG1gRNA2.3, and mIgG1gRNA2.4) (underlined, in red) in comparison with the WT sequence (mIgG1WT). Dashes: deleted nucleotides.

### Engineering of the 9E10 hybridoma cell line

To carry out the knock-in mechanism, the 1582-bp insert was composed of i) the first 18 bp of the mouse *IgG1* hinge exon, including the cysteine codon required for binding to the L chain; ii) the whole hinge region exon (105 bp) from the Antarctic fish IgM, the mouse *IgG1 CH2* and *CH3* exons; iii) a 30-bp linker; iv) mCherry coding sequence (CDS), a red fluorescent protein derived from *Discosoma sp*., as a selection marker for the correct sequence integration (Fig 1B; Fig EV4).

Compared to the mouse IgG1 hinge region, the Antarctic fish IgM hinge possesses several distinct features, such as a predominance of glycines, prolines, and asparagines (Fig EV5A and B) and the presence of two N-glycosylation sites (Fig EV5C). On the other hand, the amino acid sequence in the middle of Antarctic fish hinge recalls the core hinge of mouse IgG1 (Marquart *et al*, 1980) (Fig EV5D).

As supported by several papers, the length of homology arms represents a key element to enhance the insertion of large DNA fragments (Byrne *et al*, 2015; Howden *et al*, 2015). Thus, to optimize the efficiency and accuracy of integration, the donor construct harbored a 1005-bp 5’ and a 2421-bp 3’ homology arm, matching the sequences side of the Cas9-mediated DSB cleavage. To avoid Cas9 cutting of the donor template, the NGG PAM sequence, localized into the 5’ homology arm, was mutated into NGA.

To generate the donor plasmid, the construct was combined with 5’ and 3’ homology arms using the Gibson Assembly Protocol (see Materials and Methods section). Through the overlapping ends, the DNA fragments were cloned into the pUC19 vector, which is suitable to incorporate large inserts (Fig EV6).

To assess whether the assembly of DNA fragments had properly occurred, DNA plasmid-positive colonies were double-digested with *Hind*III and *Sph*I and analyzed on 1% agarose gel. HDR1 and HDR3 colonies showed one bright 5000-bp long band, corresponding to the donor construct, and a second one of 2600 bp, corresponding to the pUC19 plasmid. HDR2 colony showed a band of about 1500 bp, being a false positive. Overall, the correct assembly of the donor plasmid was confirmed (Fig EV7).

Next, the 9E10 hybridoma cell line was electroporated with the pX458-gRNA2 plasmid along with the donor plasmid, at a 1:1 ratio. At 24 h post-electroporation, 6.4% of GFP-positive cells were isolated by FACS (Fig EV8), relative to cells electroporated only the pX458-gRNA2 (positive control) and untreated cells (negative control), for *in vitro* expansion. Fifteen days later, 2.1% mCherry-positive cells (expressing anta-mAb) were isolated by FACS from the total GFP-positive cell population (Fig 4). Despite the high percentage of viable cells (85%), the very limited value of mCherry-positive cells is not surprising as resulted from the HDR repair.

**Figure 4.**
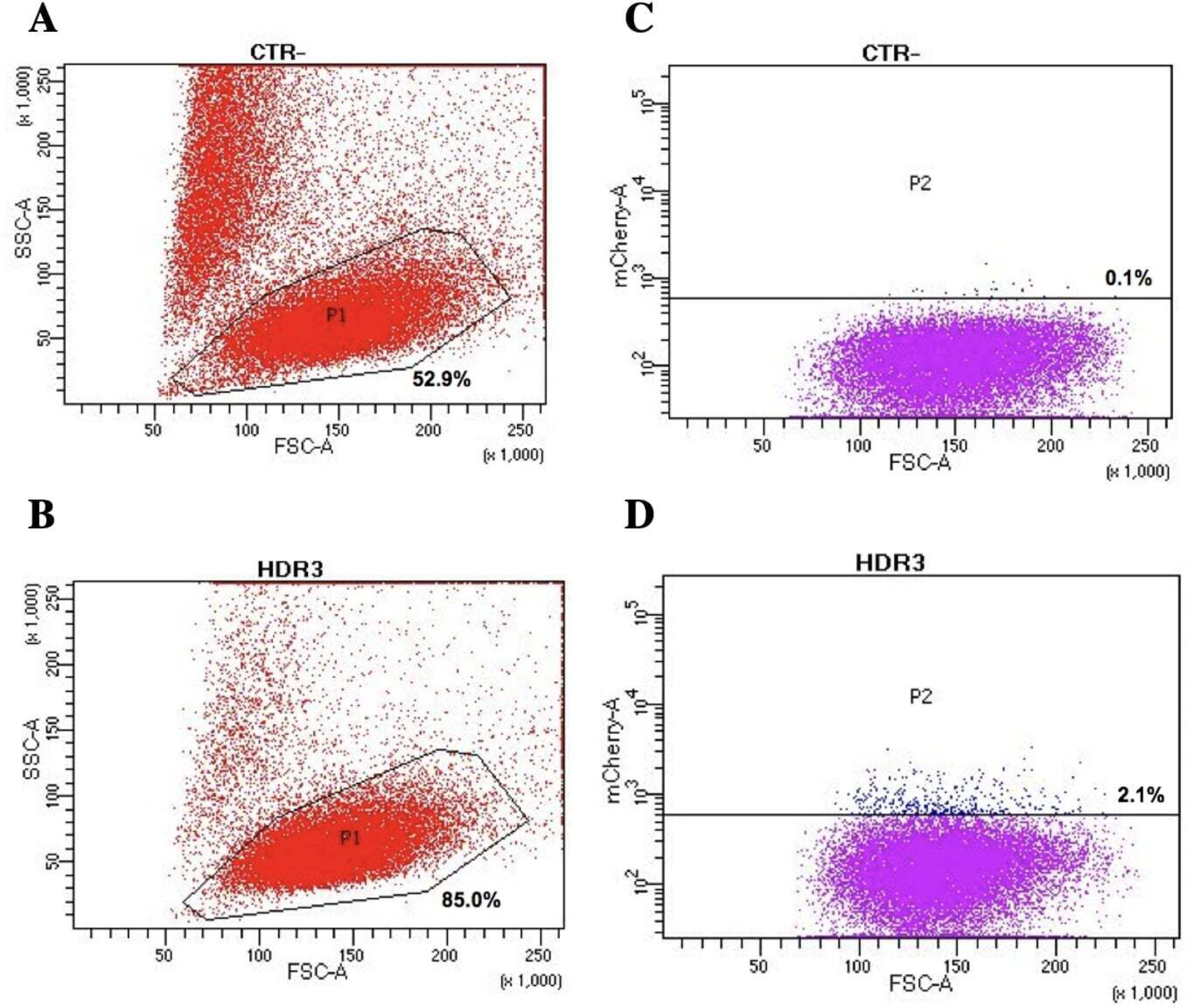
Generation of the engineered 9E10 hybridoma cell line. A–D Flow cytometry dot plots show cell viability (A and B) after electroporation with pX458, containing the gRNA2 and the donor construct, in 9E10 hybridoma cells and mCherry expression (C and D). Negative controls (A and C; CTR-) are reported.

### Biochemical and functional characterization of anta-mAbs

anta-mAbs were purified by affinity chromatography from mCherry-positive hybridoma cell supernatant. The correct production of the engineered antibody was assessed by Coomassie-Brilliant Blue-stained SDS-PAGE (Fig 5). WT mAbs showed a typical Ig pattern consisting of a 50-kDa band, corresponding to the H chain, and one of 25 kDa, corresponding to the L chain. In the case of anta-mAbs, a third band of about 75 kDa, corresponding to the expected molecular weight of the engineered H chain, was observed. (Fig 5A). Moreover, a faint band of about 150 kDa was detected for both WT and anta-mAb, more likely due to the presence of the unreduced whole antibody molecule. In the latter case, it was observed a slightly heavier band, in line with the higher molecular weight of the engineered H chain. Western blot analysis, performed with a mouse anti-mCherry as the primary antibody, validated the presence of a sharp band corresponding to the engineered H chain, carrying the mCherry protein only in anta-mAb (Fig 5B). Moreover, to clarify the nature of the additional band of 50 kDa, a control was run by omitting the primary antibody, indicating that the presence of this band was due to non-specific binding of the secondary antibody (Fig EV9). Overall, these data confirm that genome editing successfully occurred.

**Figure 5.**
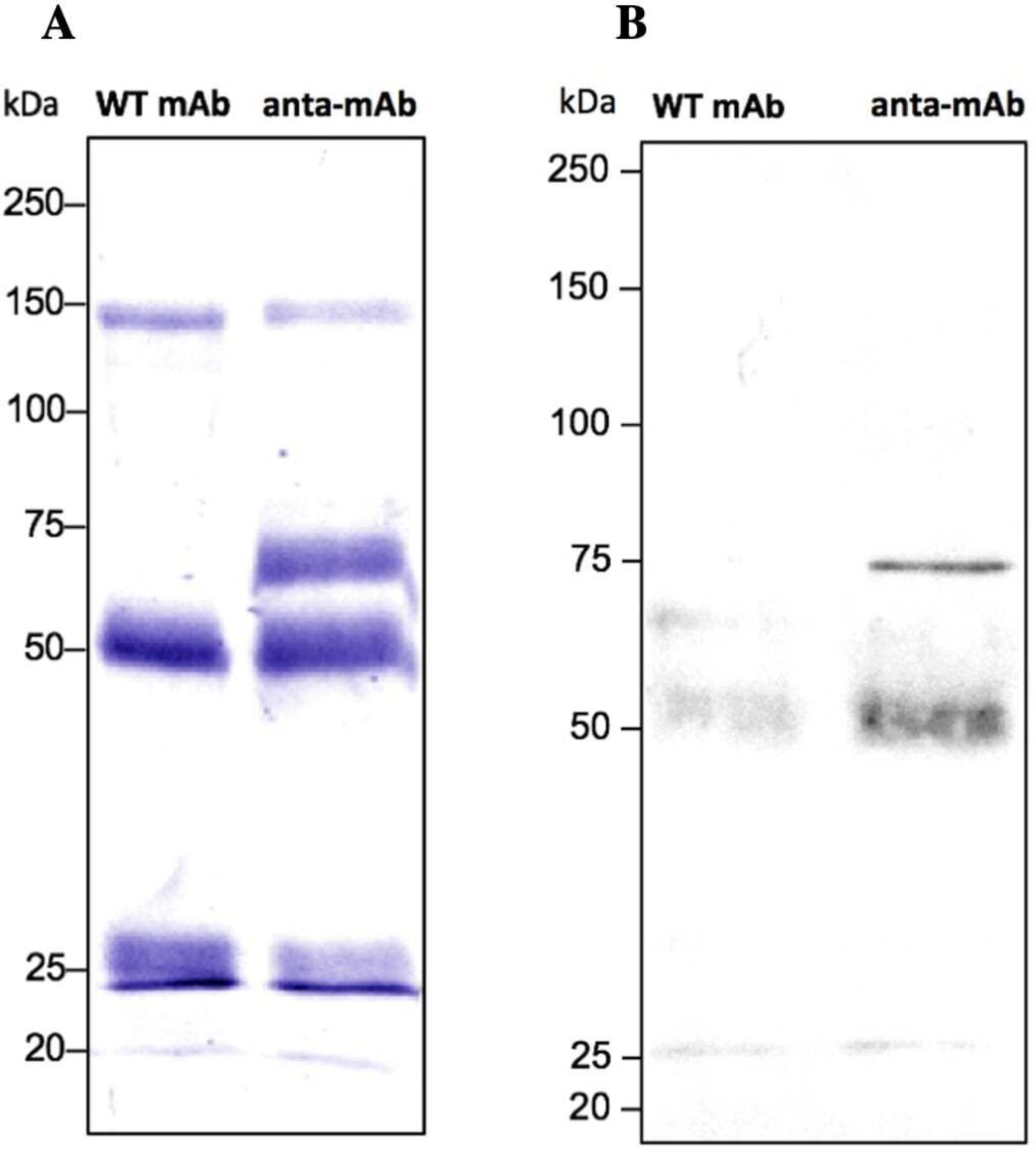
Validation of the production of anta-mAbs. A Coomassie-Brilliant Blue-stained 10% SDS-PAGE run under reducing conditions of purified WT and anta-mAbs. The H and L chain bands are marked by the arrow. B Western blot analysis was performed using a mouse anti-mCherry antibody as the primary antibody. The expected band at 75 kDa, corresponding to the engineered H chain carrying mCherry protein in anta-mAb. Molecular weight markers are shown on the left-hand side of each panel.

To test the engineered mAb affinity for its antigen (c-Myc), enzyme-linked immunosorbent assay (ELISA) was carried out using three different concentrations of c-Myc (0.12 μg/ml; 0.24 μg/ml; 0.48 μg/ml), immobilized to wells and seven serial dilutions of WT and anta-mAbs as primary antibodies (Fig 6). ELISA data confirm that mCherry-positive 9E10 hybridoma cells secreted antibodies specific for c-Myc. Moreover, anta-mAbs at the lowest dilution (1:5) were shown to readily bind the antigen at the lowest concentration (0.12 μg/ml), whereas doubling antigen concentrations (0.24 μg/ml and 0.48 μg/ml) did not result in increased binding activity (Fig 6B). In contrast to anta-mAb, it was observed for WT mAbs that increasing binding was proportional to doubling antigen concentration (Fig 6A). These results suggest that engineering of the mAb enhances the antigen sensitivity.

**Figure 6.**
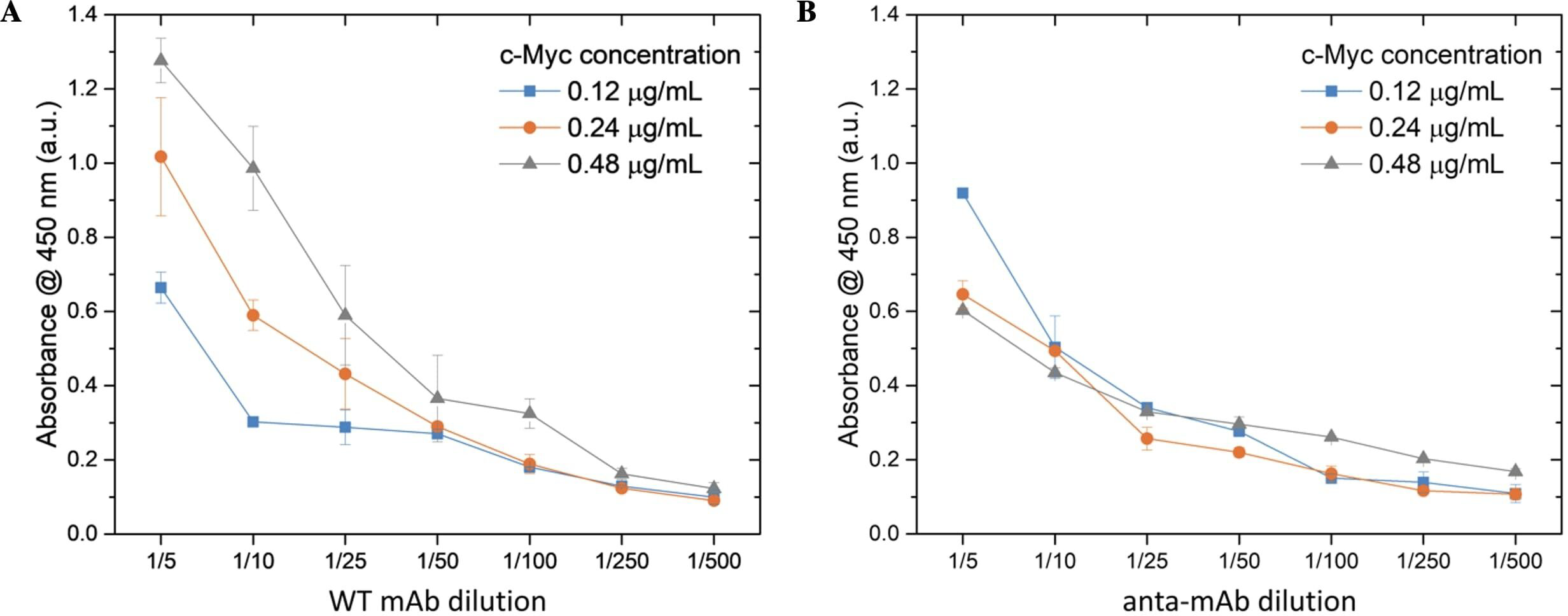
Antigen concentration-dependent binding of WT and anta-mAbs measured by Indirect ELISA. A, B Three different concentrations of antigen (purified recombinant human c-Myc protein), indicated in the upper right corner of the graph, were immobilized in wells. Serial dilutions used for mAbs are shown on the x-axis (starting from initial concentrations of 480 μg/ml). An HRP-conjugated anti-mouse IgG was used for detection. Each point is the mean of two replicate wells (*n = 2*). Error bars represent standard deviation.

Comparison between the functionality of the anta-mAb and WT mAb, was assessed, after their immobilization on a plasmonic substrate. The two mAbs underwent the same immobilization procedure on an optical transducer of plasmonic nanoislands made by an alloy of gold and silver (more information on the substrate fabrication and optical characterization in Materials and Methods section). The functionalization procedure is schematically represented in Fig 7A and described in the Materials and Methods section. Briefly, the bimetallic nanoislands (Ag/Au Nis) were capped with a MUA/MCH thiolic solution in a sufficiently high ratio (1:9) to guarantee a good spacing between mAbs molecules. Then, the carboxylic groups on MUA were activated by using EDC/NHS click chemistry to covalently bind G-Protein. The role of G-Protein is to provide good orientation of the mAbs (both wild type and engineered ones), which are recognized on their Fc region. In this way, both Fab regions are capable of recognizing their target (c-Myc). After the incubation of mAbs of both types, at the same concentration, a passivation step with BSA was performed to avoid a-specific interactions. Each functionalization step was optically monitored to measure the relative shift of the LSPR peak (Fig 7B and C) for both mAb types. The same redshifts (∼10 nm) were observed for both mAb types, while no significant redshifts were observed for the passivating step. The customized transmission setup and the typical LSP resonance of the adopted optical transducer are described in the Materials and Methods and schematized in Fig 7D. Finally, c-Myc antigen concentrations were prepared and incubated on the plasmonic substrates functionalized with the two types of mAb. The LSPR relative shifts were plotted as a function of c-Myc concentrations and fitted with the following Hill-type equation:

**Figure 7.**
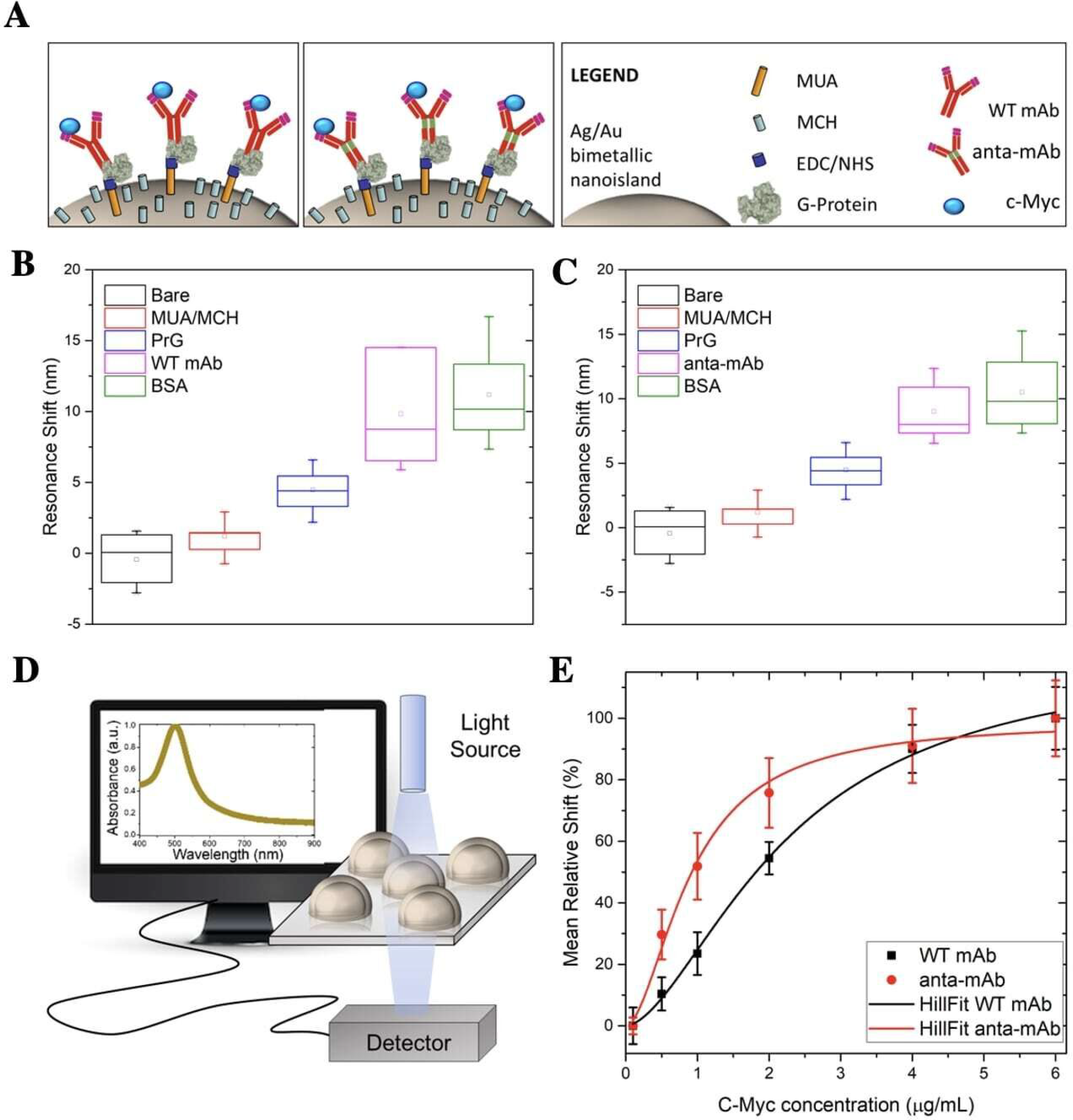
Comparison between functionalities of the WT mAb and the anta-mAb on bimetallic plasmonic nanoislands. A Schematic representation of the adopted functionalization approach. B, C Scatterplots of the measured LSPR resonance shifts of the bare bimetallic nanoislands (black) and after functionalization with MUA/MCH (red), G-Protein (PrG) (blue), WT mAb (B) and anta-mAb (C) (purple), and passivation with BSA (green) (*n ≥ 5*). D Schematic representation of the customized transmission mode setup adopted to collect the absorbance spectra of bimetallic plasmonic nanoislands. E Mean percentage relative LSP resonance shifts as a function of c-Myc concentration for WT mAb (black) and anta-mAb (red). Hill curves were used to fit the data. The vertical bars denote the standard deviation on a minimum of three independent experiments (*n ≥ 3*).

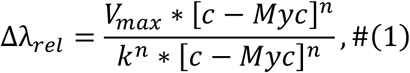

where *Δλ*_*rel*_ is the relative LSPR shift (%) calculated according to Equation 1. *V*_*max*_ is the maximum velocity of the reaction, [c-Myc] is the antigen concentration, and *n* is the Hill coefficient, providing information on the substrate binding sites. In the case of mAb molecules, we have *n = 2*, since each mAb has two Fab regions to recognize the c-Myc antigen. Finally, the mAb functionality was estimated by extracting the half-maximal concentration constant *k*. This parameter provides useful information on the apparent affinity of the mAb for the antigen. The lower the numerical value of *k*, the higher the apparent affinity for the substrate. Interestingly, the *k*-value of anta-mAb molecules resulted in 2.5-fold lower value than WT mAb, suggesting a 2.5-fold higher affinity for c-Myc antigen (Fig 7E).

## Discussion

The CRISPR/Cas9 technique has revolutionized genomic engineering and is likely one of the biggest breakthroughs for scientific research of the current century. This technique is particularly suitable and valuable for immunological studies (Cheong *et al*, 2016; Pogson *et al*, 2016; Kelton *et al*, 2017; Khoshnejad *et al*, 2018; van der Shoot *et al*, 2018) as for genetic manipulation of single cell type as well as generation of a complete gene-edited immune system, achieved in a very short time frame.

These relevant findings prompted us to employ the CRISPR/Cas9 system in the present work, aimed at generating an engineered mAb that carries a hinge region from a cold-adapted fish Ig molecule (Coscia *et al*, 2000). To generate this type of mAb, candidate gRNAs were designed to target the intronic region encompassing the *CH2* and *CH3* exons in order to avoid unintended DNA modifications in CDS during the knock-in mechanism (Doench, 2017; Hanna & Doench, 2020). Furthermore, to increase the frequency of homology-directed repair, particular attention was paid to the distance of gRNAs to the target site (Zhang *et al*, 2017). After introducing Cas9 and the gRNA into cells, we performed PCR amplification and sequencing since it has been generally considered a good approach to determine the mutation frequencies (Ran *et al*, 2013). Sequencing of PCR products revealed the same 14 bp-deletion in all gRNA1 clones and a various number of nucleotides deleted in gRNA2 clones (ranging from 6 to 10 bp). Although the limited number of sequences analyzed, this result indicates that NHEJ is the most frequently occurring mechanism for DSB repair (Jasin & Rothstein, 2013).

Next, most efforts were focused on setting up the optimal conditions to facilitate knock-in. The 5’ and 3’ flanking homology arms of the HDR donor were designed on intronic regions in order to avoid CDS knock-in (Lau *et al*, 2020). Also, the length of homology arms and the size of inserts were carefully evaluated to improve HDR efficiency (Zhang *et al*, 2017).

Biochemical analysis of purified anta-mAbs in comparison with WT mAbs confirmed that mCherry-positive hybridoma cells secreted a whole antibody molecule, as visualized by SDS-PAGE, under reducing conditions. Surprisingly, anta-mAbs showed a band of 50 kDa, corresponding to the size of the WT H chain, and one of 75 kDa, corresponding to the expected size of the engineered H chain, as confirmed by western blot analysis by using an anti-mCherry primary antibody. The presence of two differently sized H chains can be ascribed to the combination of WT alleles, NHEJ-repaired alleles, and the desired HDR-edited allele in the cell population (Pardo *et al*, 2009). Additionally, it is well described that cut-to-mutation distance influences homo or heterozygous mutation incorporation (Paquet *et al*, 2016). Thus, given that the distance of gRNA2 from the target site is about 46 bp, another possible explanation is that such distance is not short enough, the reason why more likely it promotes heterozygous editing (Paquet *et al*, 2016).

Preliminary functional data were obtained by testing WT and anta-mAbs for antigen affinity by ELISA, using a human c-Myc tag as antigen. As expected, the anta-mAb retained its antigen specificity. Surprisingly, the maximum binding was detected at the lowest antigen concentration when compared, at an equal dilution factor, to WT mAb, which performed binding with a proportional increase in antigen concentration.

ELISA data opened the possibility that the presence of the Antarctic hinge region may contribute to enhancing antigen binding. To verify this, we immobilized WT and anta-mAbs on plasmonic substrates made of highly sensitive, large-scale, bimetallic nanoislands. The functionalized substrates were then exposed to increase concentrations of the c-Myc target. The results obtained are very promising since a 2.5-fold increase in the apparent affinity for the antigen from the genetically-modified anta-mAb was observed. Since both WT and anta-mAbs are identical in the sequence of the antigen binding site, differences observed in binding activity could be ascribed to the degree of flexibility, which is mediated by different hinges. As reported in previous crystallographic studies performed on the flexibility of human Igs, the hinge-folding mode of flexibility may play a role in enhanced affinity (Roux *et al*, 1997; Roux *et al*, 1998). Length aside (22 aa vs 13 aa), features of the Antarctic fish hinge compared to those of the murine counterpart, e.g., richness in negatively charged residues and glycines, and a half number of cysteines (2 vs 4) potentially forming disulfide bonds, make this region certainly more flexible with less steric constraints. This could favor the proper orientation of antigen-binding sites on the Fab arms and, in turn, a high-affinity interaction of the anta-mAb (Adlersberg, 1976; Tan *et al*, 1990). Therefore, we attribute the increase in the apparent affinity to the mechanical mobility of the Antarctic hinge region, which can be viewed as a structural module that may contribute to local adaptive changes in flexibility. It has been hypothesized that proteins exist in an array of functional and non-functional conformational “microstates” having similar flexibilities which vary with temperature (Dong *et al*, 2018). We are, thus, confident that the hinge region, “borrowed” from a cold-adapted antibody, might properly play its role as a flexible spacer even at higher temperatures, contributing to optimal Fab orientation for antigen engagement.

Further analysis of the physicochemical properties of anta-mAb will be performed to better clarify the contribution of the Antarctic hinge region to the improvement of the binding affinity of anta-mAbs, also in terms of antigen size and antibody avidity (Oda, 2004). It will be also interesting to assess the impact of such structural modification on effector functions the engineered mAb (Vidarsson *et al*, 2014).

The results obtained from our present work could pave the way for a new class of engineered biosensors for the detection of diseases, in which not only the transducing elements are optimized, but also the biorecognition elements conferring high specificity and selectivity for a target analyte are genetically modified. In this way, biosensors could show enhanced limits of detection (LODs) and specificity for a certain target. It is well-known, indeed, that for some specific biomarkers LODs down to single-molecule level can be required. Genetically-modified mAb, therefore, could represent a promising alternative to the very expensive techniques, which are currently needed for the detection of biomarkers at very low concentrations in the body fluids (Walt *et al*, 2013; Ishii & Yanagida, 2000; Zander *et al*, 2002).

In light of these results, it is challenging to think about the development of molecular therapeutics for human diseases by using the CRISPR/Cas9 tool. To date, several studies demonstrate that Fc-engineered antibodies (Wang *et al*, 2019; Liu et al, 2020) show an increase in effector functions and serum half-life (Saunders, 2019), useful for different applications.

The results obtained from the present work have confirmed the versatility and efficiency of the CRISPR/Cas9 technology for precise genome editing and offer perspectives on further structural modifications of the antibody molecule for different purposes.

## Materials and Methods

### Hybridoma cell culture conditions

MYC 1-9E10.2 [9E10] (ATCC CRL-1729), a hybridoma cell line secreting monoclonal antibody (IgG1 subclass) against human c-Myc tag, was kept in culture in RPMI 1640 medium (Corning 10-040-CV) supplemented with 10% heat-inactivated fetal bovine serum (Corning, 35-016-CV), 100 U ml^-1^ penicillin and 100 μg/ml streptomycin (Corning, 30-002-CI). All hybridoma cells were maintained at 37 °C with 5% CO_2_ in 10 ml of culture in T-25 flasks (Falcon, 353109) and passaged every 48/72 h. All hybridoma cell lines were confirmed to be negative for *Mycoplasma* contamination (MycCellService at the Institute of Genetics and Biophysics “Adriano Buzzati-Traverso” – CNR, Naples).

### gRNAs target selection

9E10 hybridoma cell line expressing mouse anti-c-Myc monoclonal antibody (IgG1 subclass) was chosen for genome editing. Mouse IgG1 heavy chain constant region sequence was retrieved from GenBank (NCBI - accession number: AJ487681). To edit the target gene locus, candidate gRNAs were designed by using the E-CRISP Design tool (Heigwer, 2014). The gRNAs chosen showed a low rate of off-targets, expressed as the number of hits, meaning that how Cas9-cut is specific. In addition to the number of hits, other parameters were taken into consideration, such as S-Score (specificity, starting with 100), A-score (annotation, starting with zero), E-score (efficacy, is higher when sequence tends to give out of frame deletions), that are algorithms for the evaluation of gRNAs quality.

### Construction of the CRISPR plasmid containing gRNAs

pSpCas9(BB)-2A-GFP was a gift from Feng Zhang (pX458, Addgene #48138;) (Ran *et al*, 2013), used for cloning of gRNA1 and gRNA2. Two guanines were added to the 5’ end of the gRNAs to improve the U6 transcription. Both strands of the gRNAs were flanked by overhangs CACC and CAAA, respectively, for the ligation into the *Bbs*I site in the pX458 plasmid (Table EV2). DNA oligos were purchased from Eurofins Genomics Europe Sequencing GmbH and suspended to 100 μM. The top and bottom strands of gRNA1 and gRNA2 were mixed with 10X annealing buffer (1M NaCl, 100 mM Tris-HCl, pH 7.4) in a separate tube and the oligo mixtures were placed in the water bath, allowing them to cool down naturally to 30 °C to enhance annealing efficiency of oligos. The annealed oligos were diluted 1:400 in 0.5X annealing buffer.

For cloning gRNAs, pX458 plasmid was digested with *Bpi*I (*Bbs*I; Thermo, #ER1011) and gel-purified by NucleoSpin Gel and PCR Clean-up (Macherey-Nagel); gRNA oligonucleotides were ligated into pX458 plasmid with T4 DNA ligase (NEB, #M0202S). The ligation reaction was transformed into *DH5 α-*competent *E. coli*. DNA plasmid was extracted from positive colonies by using Exprep Plasmid SV mini (GeneAll) and sequenced on an ABI PRISM 3100 automated sequencer at Eurofins Genomics Europe Sequencing GmbH.

### HDR donor plasmid construction

The HDR donor plasmid was constructed following the Gibson Assembly Protocol. The CDS of the Antarctic IgM hinge region used for the construction of HDR donor plasmid was obtained from a multiple alignment of nucleotide sequences available from the 12 speciments of the Antarctic fish species *Trematomus bernacchii* (GenBank NCBI – accession number: EU884293). The closest sequence to the consensus was chosen. The insert, containing CDS encoding Antarctic hinge, mouse IgG1 CH2 and CH3 domains (GenBank NCBI – accession number: AJ487681), and mCherry protein (GenBank NCBI – accession number: AY678264), flanked by overlapping ends for Gibson Assembly Protocol, was obtained as a synthetic gene fragment (gBlocks, Integrated DNA Technologies, IDT).

PCR amplification of 5’ and 3’ homology arms, containing overlapping ends for Gibson Assembly Protocol, was performed in a final volume of 25 μl, using 2 μl of genomic DNA (20 ng), 1.25 μl of specific primers (0.5 μM), 0,5 μl dNTP mix (0.2 μM), 5 μl 5X Q5 Reaction Buffer, 0.25 μl (0.5 U) of Q5 High-Fidelity DNA Polymerase (NEB, #M0491G), up to volume with H_2_O. The following cycling conditions were used on the 5’ homology arm: 98 °C for 30 s, 30 cycles of 98 °C (10 s), 60°C (30 s), and 72 °C (30 s), with a final extension at 72 °C for 2 min. The following cycling conditions were used on the 3’ homology arm: 98 °C for 30 s, 35 cycles of 98 °C (10 s), 65.1 °C (30 s), and 72 °C (1 min), with a final extension at 72 °C for 5 min. Primers used for 5’ and 3’ homology arms amplification are reported in table S2. PCR products were analyzed on 1% agarose gel and purified by NucleoSpin® Gel and PCR Clean-up (Macherey-Nagel). 5’ and 3’ homology arms and the insert were cloned into the pUC19 vector (NEB, #N3041S) by using Gibson Assembly Master Mix (NEB, #E2611S). In 20 μl of reaction, 0.003 pmol 5’ homology arm, 0.001 pmol of 3’ homology arm, and 0.018 pmol of insert were assembled in 54.6 ng of pUC19 with 10 μl Gibson Assembly Master Mix (2X), up to volume with deionized water. The reaction was incubated at 50 °C for 1 h.

The Gibson product was transformed into *DH5 α-*competent *E. coli*. DNA plasmid was extracted from positive colonies by EndoFree Plasmid Maxi kit (QIAGEN) and double-digested with *Hind*III (NEB, #R0104S) and *Sph*I (NEB, #R0182S) at 37 °C for 1 h (Ametrano & Coscia, 2022).

### Hybridoma transfection with CRISPR/Cas9 and HDR donor plasmids

9E10 hybridoma cells were electroporated by using GenePulser Xcell Electroporation System (Bio-Rad). Cells were prepared as follows: 7 x 10^6^ were isolated and centrifuged at 1000 rpm for 4 min, washed twice with 5 ml of PBS, and centrifuged again at the same conditions. Cells were finally re-suspended in 500 μl of PBS. For the knock-out step, 5 μg of pX458 plasmid containing gRNA1 and gRNA2 were added to cells and the cell/DNA mix was transferred into a cuvette. The mixture was electroporated using the following conditions for the electroporator: 250 kV, 500 μF at the typical time constant of about 3 ms.

For the knock-in mechanism, 5 μg of pX458 with gRNA1 and 5 μg of the circular HDR donor plasmid were electroporated into cells, following the same conditions as for the knock-out step. After electroporation, cells were typically kept in culture in 1 ml of RPMI 1640 supplemented with 20% FBS in 24 well plates at a density of 200,000 cells/well (Thermo, NC-142475). After sorting, typically 24 h after electroporation, cells were recovered from 24-well plates, and progressively transferred into six-well plates (Thermo, NC-140675) and T-25 flasks (Thermo, NC-156340), following expansion.

### Flow cytometry analysis and sorting of hybridoma cells

BD Aria II FACS cell sorter (BD Biosciences) at the IBBC-IGB Facility in the Area della Ricerca CNR Napoli 1, was used for isolation of GFP and mCherry positive cells. At 24 h post-electroporation, 9E10 hybridoma cells were collected, centrifuged at 1000 rpm for 4 min, re-suspended in Sorting Buffer (PBS supplemented with 0.1% FBS), and isolated for expression of GFP and mCherry.

### Evaluation of Genome Editing in hybridoma cells

Genomic DNA of hybridoma cell lines was extracted from about 5×10^5^ cells by using PureLink™ Genomic DNA Mini Kit (Thermo, K182001). The evaluation of Cas9-mediated DSB was performed by PCR amplification by using primers that flanked the target region of gRNA1 and gRNA2 (Table EV2). The target sequence was amplified in a final volume of 25 μl using 2 μl cDNA (50 ng), 1,25 μM of specific primers (0,5 M), 0,5 μl dNTP mix (0,2 μM), 2,5 μL 10X DreamTaq Buffer (Thermo, #EP0701), 0,5 μl (1U) of DreamTaq DNA polymerase (Thermo, #EP0701), up to volume with H_2_O. The cycling parameters are as follows: 95 °C for 3 min, 35 cycles of 95 °C (30 s), 62 °C (30 s), and 72 °C (1 min), with a final extension at 72 °C for 10 min. PCR products were analyzed on 1.5% agarose gel, purified by NucleoSpin Gel and PCR Clean-up (Macherey-Nagel), and cloned into pGEM-T Easy Vector (Promega). Positive clones were identified by the blue/white screening method and sequenced on ABI PRISM 3100 automated sequencer at Eurofins Genomics Europe Sequencing GmbH.

### Production and purification of mAb from hybridoma cells

9E10 hybridoma cell lines grown in conventional medium with 10% FBS were subcultured into prewarmed Hybridoma-SFM medium (Gibco), a serum-free and very-low protein medium, suitable for monoclonal antibody production. All hybridoma cells were maintained at 37 °C with 5% CO_2_ in 10 ml of culture in T-75 flasks (Falcon) and passaged every 48/72 h. After monitoring the viable cell density using Burker Chamber Cell Counting, WT, and anta-mAb 9E10 hybridoma cell supernatants were collected every 3-5 passages. The purification of monoclonal antibodies was performed by affinity chromatography using Hitrap Protein G Column (Sigma) starting from 150 ml of hybridoma cell supernatant. After purification, the Bradford assay (AppliChem) had been performed: 2,18 mg/ml for WT mAb and 3,86 mg/ml for anta-mAb. To assess the quality of purified monoclonal antibodies, the samples were run under reducing conditions on sodium dodecyl sulfate-polyacrylamide gel electrophoresis (SDS-PAGE), 10% homogenous gel at 100 V for 3 h. Gel was stained with 0.1% Coomassie Brillant Blue R-250 (Bio-Rad), dissolved in 40% methanol and 10% acetic acid, up to volume with deionized water.

### Western blot analysis of purified mAbs

Samples were denatured with SDS-PAGE Protein Sample Buffer 2X (80 mM Tris HCl pH 6.8; 2% SDS; 10% Glycerol; 0.0006% Bromophenol blue; 2% 2-mercaptoethanol) at 95 °C for 5 min. Samples were run under reducing conditions on SDS-PAGE, 10% homogenous gel at 100 V for 3 h. Gel was transferred onto 0,2 μm nitrocellulose membrane (Protran Nitrocellulose Membranes - Whatman Schleicher & Schuell) at 100 V for 90 min. The membrane was blocked for 1 h with 5% non-fat dry milk in TBS-T 0.1% Tween-20.

For mCherry detection, the membrane was incubated with 1:1000 mouse anti-mCherry monoclonal antibody (Elabscience E-AB-20087) at 4 °C overnight, followed by 1:15000 sheep anti-mouse HRP secondary antibody (Bethyl, A90-146P) at RT for 1 h. For chemiluminescence western blot analysis of the modified antibodies, Clarity Western ECL Substrate (Bio-Rad, #1705061) was used.

### Measurement of antigen affinity by ELISA

For antigen-affinity measurements, the plates (UltraCruz ELISA Plate sc-204463) were coated with purified recombinant human c-Myc protein (Abcam ab84132), diluted in three different concentrations (0.12 μg/ml; 0.24 μg/ml; 0.48 μg/ml), in Coating Buffer at 4 °C overnight. Plates were then blocked with PBS (1X) with 1% Tween-20 and 10% BSA at RT for 2 h. After blocking, the plates underwent three washing steps with PBS supplemented with 0.5% Tween-20 (PBST). WT and anta-mAbs were then serially diluted in PBS (1X), containing 1% Tween-20 and 10% BSA, starting from the non-diluted samples, and added to the appropriate well at RT for 2 h. PBS in place of mAb and no antigen-coated wells were tested as negative controls. An HRP-conjugated anti-mouse IgG (sheep anti-mouse, Bethyl, A90-146P) had been added to the plate and incubated at RT for 1 h. ELISA detection was performed using a 1-Step Ultra TMB-ELISA Substrate Solution (Thermo, 34028) as the HRP substrate. The reaction was terminated with H_2_SO_4_ (1M). Absorbance at 450 nm was read with Gen5 All-in-One Microplate Reader (BioTek). ELISA data were analyzed with the software Gen5 V2 Data Analysis Software.

### Fabrication of bimetallic plasmonic nanoislands arrays

The fabrication of the plasmonic substrate for the mAb functionality assessment was performed by re-adapting the protocols (Qiu *et al*, 2018; Bhalla *et al*, 2018; Miranda *et al*, 2020) in a class 1000 cleanroom. Briefly, glass coverslips (24 mm x 24 mm) were sonicated in acetone and isopropanol for 2 min, respectively, and dried under a nitrogen stream. A polydimethylsiloxane (PDMS) shadow mask with 8 mm diameter holes was placed on the top of coverslips to achieve the formation of nanostructures in controlled areas. Then, the coverslips, covered by the shadow mask, were placed in a high-vacuum chamber of a thermal evaporator to undergo thermal evaporation. Once a vacuum pressure of 10^-6^ mbar was achieved, Ag thin film (final thickness of 2 nm) was deposited on the samples with a deposition rate of 0.2 Å/s. Then, after the restoration of the vacuum pressure, another deposition process of Au (final thickness of 2 nm) was performed at the same deposition rate. The samples were brought to atmospheric pressure and removed from the thermal evaporator chamber. Immediately, they were placed into a furnace for rapid thermal annealing and dewetted at 560 °C for 3 h to obtain bimetallic nanoislands.

### Optical characterization of bimetallic plasmonic nanoislands arrays

The optical absorbance spectra of the bimetallic plasmonic arrays were recorded using a customized transmission optical setup (Miranda *et al*, 2021; Miranda *et al*, 2022). Briefly, a halogen light source was conveyed to the sample by a Thorlabs optical fiber with a collimator at its end. The light transmitted by the sample was collected by another optical fiber connected to a spectrometer (Filmetrics 2020). The transmitted spectra (400-900 nm) were analyzed by OriginPro 8 Free Version and the peak analysis was performed to measure the LSPR position.

### Surface functionalization of bimetallic plasmonic nanoislands arrays

A PDMS well (10 mm diameter) was attached to the 8 mm plasmonic bimetallic transducers to keep constant volumes during the functionalization procedure. The PDMS well was designed to safely contain 500 μl reaction volume for all the incubation steps. First, the samples were exposed to a 1:1 mixture of 1 mM 11-mercaptoundecanoic acid (MUA) and 9 mM 6-mercapto hexanol (MCH) in ethanol (95%) overnight (16-18 h) at 4 °C in a humid chamber to prevent ethanol evaporation. After the incubation, the samples were rinsed with ethanol and then with 1 mM MCH to saturate the Ag/Au nanoislands surface. This step was followed by three washing steps in ethanol and drying under a nitrogen stream. For the activation of carboxyl acid groups of MUA, the bimetallic nanoislands were incubated with a solution of EDC (40 mM) and NHS (16 mM) in MES buffer for 30 min at RT. Then, the samples were incubated with a PBS (1X) solution of G-Protein (200 μg/ml) for 2 h at 4 °C. After three washing steps (one in PBS and two in MilliQ water), the functionalization procedure was completed by splitting the samples into two groups. The first group was incubated with WT mAb (PBS 1X, 10 μg/mL), while the second one was incubated with anta-mAb (PBS 1X, 10 μg/ml) for 2 h at 4 °C. Each functionalization step was monitored by recording the absorbance spectra and measuring the relative variation of the LSPR peak with respect to the initial position. The functionalization was performed on a minimum of three samples for each mAb type (*n ≥ 3*). Before incubation with target BSA solution (0.058 mg/ml, PBS 1X) was used to passivate the plasmonic transducer surface. Finally, the binding constants of the two mAb types were assessed by monitoring the relative LSPR shift upon incubation of the functionalized substrates to increasing concentrations (from 0.5 to 6 μg/ml) of c-Myc (PBS 1X), each incubated on the samples for 2 h at 4 °C. The percentage relative shifts (*Δλ*_*rel*_) measured as

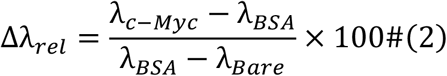

were fitted by a Hill-type curve from which the affinity constants of the two mAb types for the c-Myc antigen were estimated (*n ≥ 3*). In Equation 1, *λ*_*c-Myc*_, *λ*_*BSA*_, and *λ*_*Bare*_ denote the measured LSPR wavelengths after incubation with the c-Myc target, BSA passivation step, and before the functionalization, respectively.

## Data and materials availability

All data are available in the main text or the supplementary materials. The described plasmids used in this study are deposited in plasmid repository of Addgene (www.addgene.org/). Additional data and materials related to this paper may be requested from the authors.

## Acknowledgments

We are grateful to Luciana D’Apice (Institute of Biochemistry and Cell Biology, National Research Council of Italy, Naples, Italy) for useful discussion and suggestions.

The work was supported by CNR@Projects SAC.AD002.173.026 TIPPS (Tracking and Identification of asymptomatic Patients through engineered antibodies bioconjugated plasmonics in a Pandemic Scenario).

## Author contributions

**Alessia Ametrano:** Conceptualization; formal analysis; validation; investigation; methodology; writing – original draft; writing – review and editing. **Bruno Miranda:** Formal analysis; validation; investigation; methodology; writing – original draft; writing – review and editing. **Rosalba Moretta:** Validation; investigation; methodology. **Principia Dardano:** Validation; investigation; methodology. **Luca De Stefano:** Funding acquisition; resources; validation; investigation; methodology; writing – original draft; project administration; writing – review and editing. **Umberto Oreste:** Formal analysis; writing – review and editing. **Maria Rosaria Coscia:** Conceptualization; resources; formal analysis; supervision; funding acquisition; methodology; writing – original draft; project administration; writing – review and editing.

## Tables

**Table EV1.**
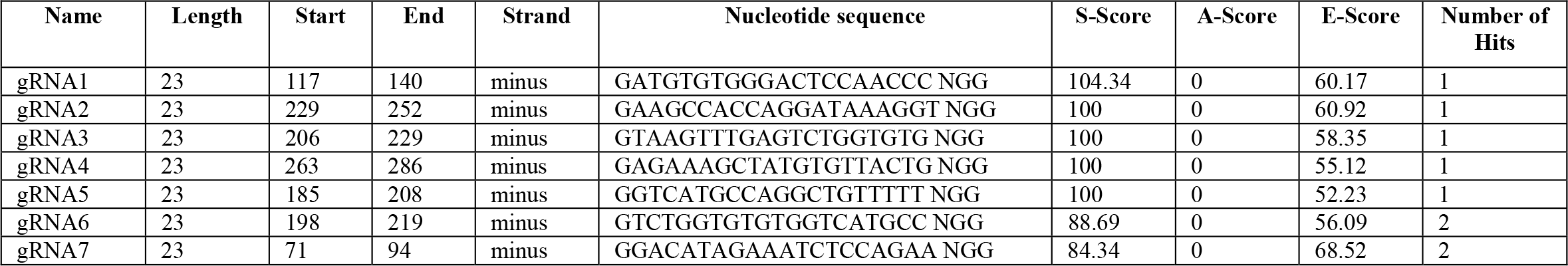
Design of gRNAs for CRISPR/Cas9 Genome Editing by the bioinformatic tool E-CRISP.

**Table EV2.**
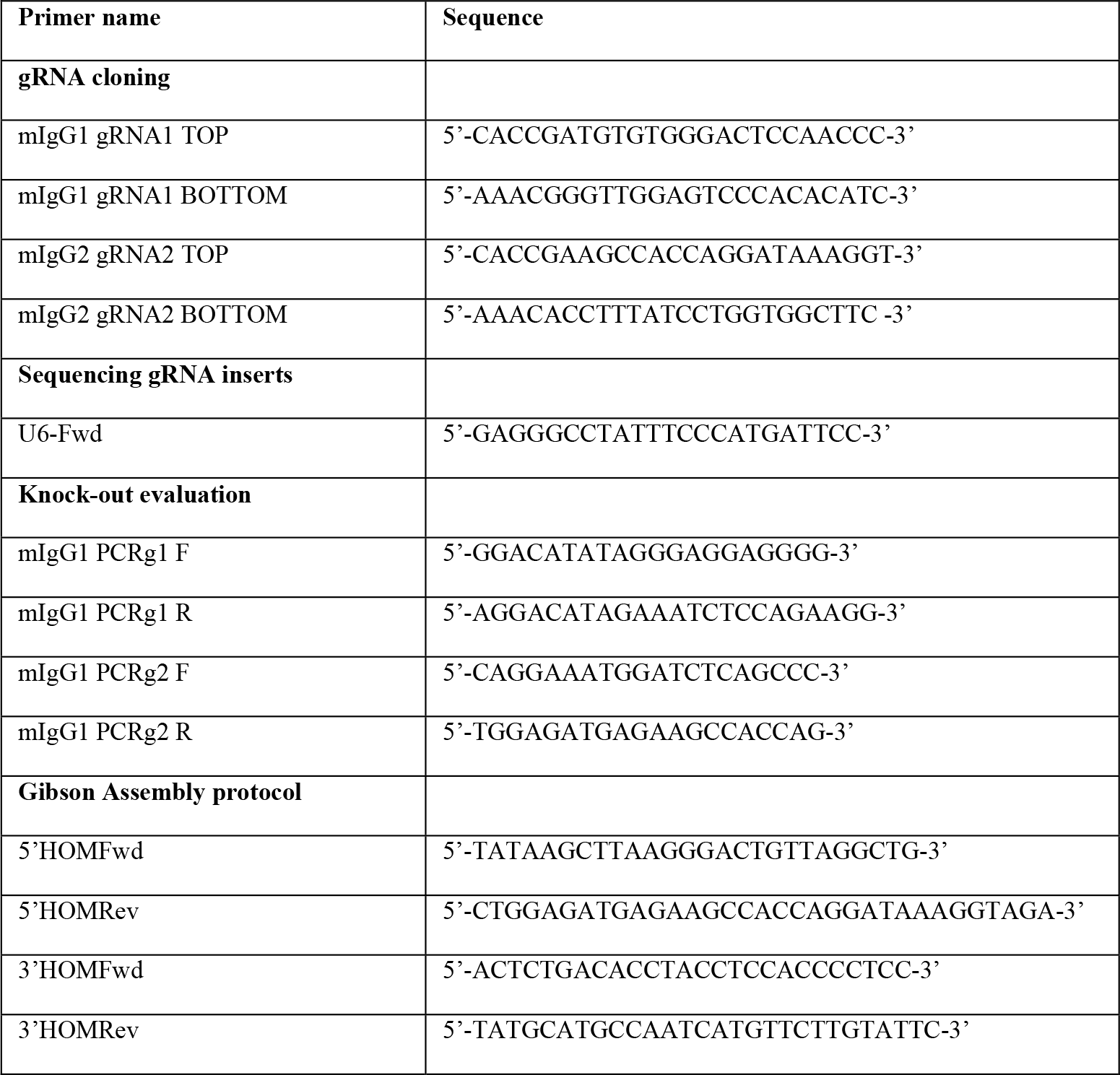
List of primers used in different experiments.

## Expanded View legends

**Figure EV1. Annotated sequence of mouse IgG1 heavy chain constant region gene (partial CDS)**.

The intronic sequence between the first constant (CH1) and the hinge region exon (highlighted in grey) represents the target region of the two gRNAs. Accession number: AJ487681.

**Figure EV2. Schematic representation for cloning gRNA oligonucleotides into the CRISPR plasmid pX458**.

The gRNA1 (highlighted in green) and gRNA2 (highlighted in light blue) oligos contain overhangs for ligation into the pair of *Bbs*I sites (in red) in pX458, with the top and bottom strand orientations matching the sequence of mouse IgG1 heavy chain constant region gene.

**Figure EV3. Evaluation of gRNAs cloning in the CRISPR plasmid pX458**.

Representative nucleotide sequences of the CRISPR plasmids pX458 containing gRNA1 (bold, highlighted in green) and gRNA2 (bold, highlighted in cyan), respectively.

**Figure EV4. Deduced amino acid sequence of the donor construct**.

**Figure EV5. Characterization of *Trematomus bernacchii* IgM hinge region in comparison with that of mouse IgG1**.

A, B Amino acid composition of (A) mouse IgG1 and (B) *T. bernacchii* hinge region.

C Prediction of N-glycosilation sites of *T. bernacchii* hinge region.

D Alignment of deduced amino acid sequence of mouse IgG1 and *T. bernacchii* IgM hinge region. Conserved cysteines are reported in bold. Below the alignment, identical amino acid residues are marked with an asterisk, positions where only one sequence shows a different amino acid residue are marked with a dot.

**Figure EV6. Nucleotide sequences of the three DNA components of the donor construct**.

Insert, 5’ and 3’ homology arms, with the respective overlapping ends (white, bold, highlighted in magenta and brown), required for Gibson Assembly Protocol. *Hind*III (bold, yellow) and *Sph*I (bold, red) sites for cloning into the pUC19 plasmid are also reported.

**Figure EV7. Agarose gel electrophoresis of double-digested DNA plasmid colonies by *Hind*III and *Sph*I**.

The green arrow points the band corresponding to the donor construct, whereas the blue arrow points the band corresponding to the pUC19 vector, used as negative control.

**Figure EV8. Generation of engineered 9E10 hybridoma cell line**.

A–F Flow cytometry dot plots show cell viability following electroporation protocol (A– C) and the Cas9 expression (D–F) in 9E10 hybridoma cells after electroporation with pX458 containing gRNA2 and HDR3 donor plasmid. Negative (A and D) and positive (B and E) controls are reported. Data were collected 15 days before sorting for mCherry expression.

**Figure EV9. Control of anti-mCherry antibody specificity**.

Purified WT and anta-mAbs were separated by 10% SDS-PAGE and transferred onto a nitrocellulose membrane. Western blot analysis was performed by omitting the primary antibody.

## References

Adlersberg JB (1976) The immunoglobulin hinge (interdomain) region. Ric Clin Lab 6: 191– 205

Ametrano A, Coscia MR (2022) Production of a chimeric mouse-fish monoclonal antibody by the CRISPR/Cas9 technology. Methods Mol Biol 2498: 337–350

Ashoor DN, Ben Khalaf N., Bourguiba-Hachemi S, Marzouq MH, Fathallah MD (2018) Engineering of the upper hinge region of human IgG1 Fc enhances the binding affinity to FcγIIIa (CD16a) receptor isoform. Protein Eng Des Sel 31: 205–212

Baker MD, Pennell N, Bosnoyan L, Shulman MJ (1988) Homologous recombination can restore normal immunoglobulin production in a mutant hybridoma cell-line. Proc Natl Acad Sci USA 85: 6432–6436

Bhalla N, Sathish S, Galvin CJ, Campbell RA, Sinha A, Shen AQ (2018) Plasma-assisted large-scale nanoassembly of metal-insulator bioplasmonic mushrooms. ACS Appl Mater Interfaces 10: 219–226

Byrne SM, Ortiz L, Mali P, Aach J, Church GM (2015) Multi-kilobase homozygous targeted gene replacement in human induced pluripotent stem cells. Nucleic Acids Res 43: e21

Carayannopoulos L, Capra JD (1993) Immunoglobulins, structure and function. Fundamental Immunology. Raven Press, New York, USA

Cheong T-C, Compagno M, Chiarle R 2016 Editing of mouse and human immunoglobulin genes by CRISPR/Cas9 system. Nat Commun 7: 1–10

Cong L, Ran FA, Cox D, Lin S, Barretto R, Habib N, Hsu PD, Wu X, Jiang W, Marraffini LA, Zhang (2013) Multiplex genome engineering using CRISPR/Cas systems. Science 339: 819– 823

Coscia MR, Morea V, Tramontano A, Oreste U (2000) Analysis of a cDNA sequence encoding the immunoglobulin heavy chain of the Antarctic teleost Trematomus bernacchii. Fish Shellfish Immunol 10: 343–357

Coscia MR, Varriale S, Giacomelli S, Oreste U (2011) Antarctic teleost immunoglobulins: more extreme, more interesting. Fish Shellfish Immunol 31: 688–696

Coscia MR, Giacomelli S, Oreste U (2012) Allelic polymorphism of immunoglobulin heavy chain genes in the Antarctic teleost Trematomus bernacchii. Mar Genom 8: 43–48

Dell’Acqua WF, Cook KE, Damschroder MM, Woods RM, Wu H (2006) Modulation of the effector functions of a human IgG1 through engineering of its hinge region. J Immunol 177: 1129–1138

Doench JG, Hartenian E, Graham DB, Tothova Z, Hegde M, Smith I, Sullender M, Ebert BL, Xavier RJ, Root DE (2014) Rational design of highly active sgRNAs for CRISPR-Cas9-mediated gene inactivation. Nat Biotechnol 32: 1262–1267

Doench JG (2017) Am I ready for CRISPR? A user’s guide to genetic screens. Nat Rev Genet 19: 67–80

Dong Y, Liao M, Meng X, Somero GN (2018) Structural flexibility and protein adaptation to temperature: molecular dynamics analysis of malate dehydrogenases of marine molluscs. Proc Natl Acad Sci USA 115: 1274–1279

Doudna JA, Charpentier E (2014) The new frontier of genome engineering with CRISPR-Cas9. Science 346:1258096

Doudna JA, Sontheimer EJ (2014) The use of CRISPR/Cas9, ZFNs, and TALENs in generating site-specific genome alterations. Methods Enzymol 546

Fell HP, Yarnold S, Hellström I, Hellström KE, Folger KR (1989) Homologous recombination in hybridoma cells - heavy-chain chimeric antibody produced by gene targeting. Proc Natl Acad Sci USA 86: 8507–8511

Frenzel A, Hust M, Schirrmann T (2013) Expression of recombinant antibodies. Front Immunol 4: 217

Hanna RE, Doench JG (2020) Design and analysis of CRISPR–Cas experiments. Nat Biotechnol 38: 813–823

Heigwer F, Kerr G, Boutros M (2014) E-CRISP: Fast CRISPR target site identification. Nat Methods 11: 122–123

Hermans P, van Soolingen D, Bik EM, de Haas PE, Dale JW, van Embden JD (1991) Insertion element IS987 from Mycobacterium bovis BCG is located in a hot-spot integration region for insertion elements in Mycobacterium tuberculosis complex strains. Infect Immun 59: 2695– 2705

Ho BK, Coutsias EA, Seok C, Dill KA (2005) The flexibility in the proline ring couples to the protein backbone. Protein Sci 14: 1011–1018

Ishii Y, Yanagida T (2000) Single molecule detection in life sciences. Single Mol 1: 5–16

Ishino Y, Shinagawa H, Makino K, Amemura M, Nakata A (1987) Nucleotide sequence of the Iap gene, responsible for alkaline phosphatase isozyme conversion in Escherichia coli, and identification of the gene product. J Bacteriol 169: 5429–5433

Jasin M, Rothstein R (2013) Repair of strand breaks by homologous recombination. Cold Spring Harb Perspect Biol 5: 11

Jin, S. Sun Y, Liang X, Gu X, Ning J, Xu Y, Chen S, Pan L (2022) Emerging new therapeutic antibody derivatives for cancer treatment. Signal Transduct Target Ther 7: 39

Kelton W, Waindok AC, Pesch T, Pogson M, Ford K, Parola C, Reddy ST (2017) Reprogramming MHC specificity by CRISPR-Cas9-assisted cassette exchange. Sci Rep 7: 457752017

Khoshnejad M, Brenner JS, Motley W, Parhiz H, Greineder CF, Villa CH, Marcos-Contreras OA, Tsourkas A, Muzykantov VR (2018) Molecular engineering of antibodies for site-specific covalent conjugation using CRISPR/Cas9. Sci Rep 8: 1760

Köhler G, Milstein C (1975) Continuous cultures of fused cells secreting antibody of predefined specificity. Nature 256: 495–497

Lau CH, Tin C, Suh Y. (2020) CRISPR-based strategies for targeted transgene knock-in and gene correction. Fac Rev 9: 20

Liu L (2020) Potent neutralizing antibodies against multiple epitopes on SARS-CoV-2 spike. Nature 584: 450–456

Liu R, ldham RJ, Teal E, Beers SA, Cragg MS (2020) Fc-engineering for modulated effector functions-improving antibodies for cancer treatment. Antibodies 9: 64

Mao Z, Bozzella M, Seluanov A, Gorbunova V (2008) DNA repair by nonhomologous end joining and homologous recombination during cell cycle in human cells. Cell Cycle 7: 2902– 2906

Marquart M, Deisenhofer J, Huber R, Palm W (1980) Crystallographic refinement and atomic models of the intact immunoglobulin molecule Kol and its antigen-binding fragment at 3.0 Å and 1.9 Å resolution. J Mol Biol 141: 369–391

Howden SE, Maufort JP, Duffin BM, Elefanty AG, Stanley EG, Thomson JA (2015) Simultaneous reprogramming and gene correction of patient fibroblasts. Stem Cell Rep. 5: 1109–1118

Miranda B, Chu K-Y, Maffettone PL, Shen AQ, Funari R (2020) Metal-enhanced fluorescence immunosensor based on plasmonic arrays of gold nanoislands on an etched glass substrate. ACS Appl. Nano Mater 10: 10470

Miranda B, Moretta R, De Martino S, Dardano P, Rea I, Forestiere C, De Stefano L (2021) A PEGDA hydrogel nanocomposite to improve gold nanoparticles stability for novel plasmonic sensing platforms. J Appl Phys 129: 033101

Miranda B, Moretta R, Dardano P, Rea I, Forestiere C, De Stefano L (2022) H3 (hydrogel-based, high-sensitivity, hybrid) plasmonic transducers for biomolecular interactions monitoring. Adv Mater Technol 7: 2101425

Mojica FJM, Montoliu L (2016) On the origin of CRISPR-Cas technology: from prokaryotes to mammals. Trends Microbiol 24: 811–820

Oda M (2004) Antibody flexibility observed in antigen binding and its subsequent signaling. J Biol Macromol 4: 45–56

Paquet D, Kwart D, Chen A, Sproul A, Jacob S, Teo S, Olsen KM, Gregg A, Noggle S, Tessier-Lavigne M (2016) Efficient introduction of specific homozygous and heterozygous mutations using CRISPR/Cas9. Nature 533: 125–129

Pardo B, Gómez-González B, Aguilera A (2009) DNA repair in mammalian cells: DNA double-strand break repair: how to fix a broken relationship. Cell Mol Life Sci 66: 1039–1056

Pogson M, Parola C, Kelton WJ, Heuberger P, Reddy ST (2016) Immunogenomic engineering of a plug-and-(dis)Play hybridoma platform. Nat Commun 7: 12535

Qiu G, Ng SP, Wu C-ML (2018) Bimetallic Au-Ag alloy nanoislands for highly sensitive localized surface plasmon resonance biosensing. Sens Actuators B: Chem 265: 459–467

Ran FA, Hsu PD, Wright J, Agarwala V, Scott DA, Zhang F (2013) Genome engineering using the CRISPR-Cas9 system. Nat Prot 8: 2281–2308

Roux KH, Strelets L, Michaelsen TE (1997) Flexibility of human IgG subclasses. J Immunol 159: 3372–3382

Roux KH, Strelets L, Brekke OH, Sandlie I, Michaelsen TE (1998) Comparisons of the ability of human IgG3 hinge mutants, IgM, IgE, and IgA2, to form small immune complexes: a role for flexibility and geometry. J Immunol 161: 4083–4090

Saunders KO (2019) Conceptual approaches to modulating antibody effector functions and circulation half-life. Front Immunol 10: 1296

Suzuki S, Annaka H, Konno S, Kumagai I, Asano R (2018) Engineering the hinge region of human IgG1 Fc-fused bispecific antibodies to improve fragmentation resistance. Sci Rep 8: 17253

Tan LK, Shopes RJ, Oi V, Morrison SL (1990) Influence of the hinge region on complement activation, C1q binding, and segmental flexibility in chimeric human immunoglobulins. Proc Natl Acad Sci USA 87: 162–166

Urnov FD, Miller JC, Lee YL, Beausejour CM, Rock JM, Augustus S, Jamieson AC, Porteus MH, Gregory PD, Holmes MC (2005) Highly efficient endogenous human gene correction using designed zinc-finger nucleases. Nature 435: 646–651

van der Shoot JMS, Fennemann FL, Valente M, Dolen Y, Hagemans IM, Becker AMD, Le Gall CM, van Dalen D, Cevirgel A, van Bruggen JAC et al (2019) Functional diversification of hybridoma-produced antibodies by CRISPR/HDR genomic engineering. Sci Adv 5: eaaw1822

Vidarsson G, Dekkers G, Rispens T (2014) IgG subclasses and allotypes: from structure to effector functions. Front Immunol 5: 520

Walt DR (2013) Optical methods for single molecule detection and analysis. Anal Chem 85: 1258–1263

Wang S (2018) Advances in the production of human monoclonal antibodies. Antib Technol J 1: 1–4

Wang L -X, Tong X, Li C, Giddens JP, Li T (2019) Glycoengineering of antibodies for modulating functions. Annu Rev Biochem 88: 433–459

Woolf TM (1998) Therapeutic repair of mutated nucleic acid sequences. Nat Biotechnol 16: 341–344

Xu HT, Xiao CH, Chen W, Li CA, Meyer Q, Wu D, Wu L, Cong F, Zhang J, Liu S et al (2015) Sequence determinants of improved CRISPR sgRNA design. Genome Res 25: 1147– 1157

Yan B, Boyd D, Kaschak T, Tsukuda J, Shen A, Lin Y, Chung S, Gupta P, Kamath A, Wong A, Vernes JM et al (2012) Engineering upper hinge improves stability and effector function of a human IgG1. J Biol Chem 287: 5891–5897

Zander C, Enderlein J, Keller RA (2002) Single molecule detection in solution: methods and applications, Wiley-VCH

Zhang JP, Li XL, Li GH, Chen W, Arakaki C, Botimer GD, Baylink D, Zhang L, Wen W, Fu YW et al (2017) Efficient precise knockin with a double cut HDR donor after CRISPR/Cas9-mediated double-stranded DNA cleavage. Genome Biol 18: 35

